# A β-secretase modulator decreases Tau pathology and preserves short-term memory in a mouse model of neurofibrillary degeneration

**DOI:** 10.1101/2021.04.02.438175

**Authors:** Marie Tautou, Sabiha Eddarkaoui, Florian Descamps, Paul-Emmanuel Larchanché, Mélanie Dumoulin, Chloé Lamarre, David Blum, Luc Buée, Patricia Melnyk, Nicolas Sergeant

## Abstract

A structure-activity relationship has enabled us to identify two molecules, MAGS02-14 and PEL24-199, sharing a β-secretase modulatory effect but having or not a lysosomotropic activity, respectively. More importantly, MAGS02-14 and PEL24-199 only differ from each other by a single nitrogen atom. However, which of the lysosomotropic and/or β-secretase modulating activities is necessary for the pharmacological effect *in vivo* remains ill-defined. To address this question, the THY-Tau22 transgenic model of NFD was treated for 6 weeks in a curative paradigm and short-term memory, Tau burden, and inflammatory processes were studied. PEL24-199, possessing only the β-secretase modulatory activity, was shown to restore the short-term memory and to reduce NFD. This effect was associated with reduced phosphorylation of Tau, increased phosphatase expression, and a decrease of astrogliosis. Our results therefore suggest that the lysosomotropic activity may be dispensable for the effect on both Aβ and Tau pathologies.

## 1 Introduction

Alzheimer’s disease (AD) is a neurodegenerative disease defined by the presence of two neuropathological lesions: intraneuronal aggregates of Tau proteins and extracellular accumulation of toxic Aβ peptides respectively referred to as Tau and amyloid pathologies. Aβ peptides are generated by sequential cleavages of the Amyloid Precursor Protein (APP). The β-secretase (BACE1) endoprotease catalyzes the first cleavage of the amyloidogenic pathway, generating Aβ peptides after a second γ-secretase endoproteolytic cleavage (for a review see Müller et al., 2017). Tau pathology corresponds to the progressive accumulation and aggregation of abnormally and hyperphosphorylated isoforms of the microtubule-associated protein Tau, *in fine* forming the so-called neurofibrillary tangles (NFT) (Buee et al., 2000; Liu et al., 2012; Gao et al., 2018). Lesions and cognitive impairment are primary specific criteria for the definition and diagnosis of AD, suggesting that the pathophysiological processes underlying the development of these lesions are tightly linked to the disease and distinguishes AD from other neurodegenerative diseases. Most of the 121 agents currently in the AD-drug development pipeline are disease-modifying therapies against either Aβ or Tau, however separately (Cummings et al., 2020).

An accumulating body of evidence suggests that APP metabolism regulates Tau expression, notably via the inhibition of β-secretase that reduces intracellular Tau protein (Moore et al., 2015). Notably, at the crossroads of APP and Tau metabolisms are the cellular protein homeostasis systems: autophagy and the endosome/lysosome pathways. These degradation systems play a central role in removing misfolded or abnormal cellular components (Frake et al., 2015). Perturbed trafficking of lysosomal vesicles and enzymes including the intravesicular accumulation of substrates is characteristic of lysosomal storage disorders. Such lysosomal system dysfunctions have been reported in AD and in several Tauopathies suggesting the dysfunction of proteostasis systems (Nixon and Yang, 2011; Piras et al., 2016). The autophagic flow leading to autophagosome formation through the fusion of autophagosomes with lysosomes is a key process that can be blocked by lysosomotropic agents such as chloroquine (Tam et al., 2014; Mauthe et al., 2018).

We previously described molecules having a chloroquinoline nucleus substituted with an N, N’-disubstituted piperazine moiety. These family of molecules are acting on the autophagic/endolysosomal systems some of which were shown to be effective against both amyloid and Tau pathologies *in vitro* and *in vivo* (Melnyk et al., 2015; Sergeant et al., 2019). A ligand-based approach allowed us to define a pharmacophore and synthesize multiple compounds with different scaffolds derived from this pharmacophore (Gay et al., 2018). Among these new compounds, two differ only by one nitrogen atom for MAGS02-14 (compound 30 in Gay et al., 2018) substituted by a carbon atom at the same position for PEL24-199 (compound 31 in Gay et al., 2018). Although having a different chemical structure, MAGS02-14 exhibits a lysosomotropic activity comparable to chloroquine and a β-secretase non-competitive inhibitory activity. In contrast, PEL24-199 only had the non-competitive β-secretase inhibitory activity with a strongly reduced lysosomotropic activity. Indeed, MAGS02-14 treated cells were exhibiting swelling intracellular vesicles and an accumulation of LC3 and p62 markers indicative of an inhibition of the autophagy flux. Expression and localization of these markers were not modified by PEL24-199 treatment while Aβ 1-40 / 1-42 production was repressed both by MAGS02-14 and PEL24-199 (Gay et al., 2018). This inhibition of the autophagic flux for MAGS02-14 and no lysosomotropic activity for PEL24-199 was then associated with a shared β-secretase modulatory effect.

Modulation of APP metabolism using either β- or γ-secretase inhibitors regulates the dosage of Tau protein in human-derived cerebral cortical neurons (Moore et al., 2015). Moreover, we previously showed that molecules used for the scaffold design of both MAGS02-14 and PEL24-199 were efficient to reduce both amyloid and tau pathologies *in vivo* in a preventive paradigm (Sergeant et al., 2019). Herein, we investigated if either MAGS02-14 and/or PEL24-199 could be efficient on Tau pathology *in vivo* in a mouse model of NFD and which among the lysosomotropic or β-secretase modulatory activity is necessary for the improvement of the cognitive function and the associated Tau pathology.

## 2 Material and Methods

### Animals

In this study, we used females THY-Tau22 transgenic and wild type (WT) littermates (C57Bl/6J genetic background), obtained by crossing THY-Tau22 heterozygous males C57Bl/6J with WT females. All animals were housed in a pathogen-free facility in a 12/12-hour light-dark cycle and maintained under a constant temperature of 22°C, at 5 to 6 animals per cage (Techniplast Cages 1284L), with *ad libitum* access to food and water. The animals were used in compliance with European standards for the care and use of laboratory animals and experimentations conducted in this study were authorized by the French Direction of Veterinary Services with the approved registration number APAFIS#10392-201706231206250v4.

### Drug treatments

PEL24-199 and MAGS02-14 compounds were synthesized according to our previously described procedure (Gay et al., 2018). A safety pilot study was performed in WT animals treated for one month to establish the innocuousness of compounds MAGS02-14 and PEL24-199 at a dose of 1mg/kg and 5mg/kg. For the present study, animals were randomly distributed and THY-Tau22 and WT mice were treated for 6 weeks, starting at 6 months of age. MAGS02-14 or PEL24-199 treatment was delivered in the drinking water at a final concentration of 1 mg/kg, i.e. 12.5 μg/mL for drinking solutions considering an average weight of 25 g/mouse drinking 4 mL per day. Drinking bottles were changed once per week as aqueous solutions of compounds MAGS02-14 and PEL24-199 were previously demonstrated to be stable during more than one week. The volume of solution consumed by the mice was measured throughout the treatment period.

### Behavioral tests

#### Anxiety

Anxiety, which could interfere with a memory test, was evaluated in treated and untreated animals by the Elevated plus maze (EPM). Mice were placed in the center of a plus-shaped maze consisting of two 10-cm-wide open arms and two 10-cm-wide enclosed arms elevated at 50 cm above the floor. Parameters including distance moved, velocity, as well as the number of entries into each arm, time spent in the open versus the closed arms, and percentage of open arms entries, were measured during 5 minutes and acquired by video recording using EthoVision video tracking equipment and software (Noldus Information Technology, Paris, France) in a dedicated room.

#### Short-term spatial memory

Short-term spatial memory was assessed using the Y-maze task. The Y-maze task consists of three 10-cm-wide enclosed arms surrounded by four spatial clues. One of the two arms opposite to the starting (S) arm was alternatively closed during the learning phase. Each mouse was positioned in the starting arm and was free to explore the maze for 5 min. Then during the retention phase of 2 min, the mouse was returned to the home cage. During the test phase of 5 min, the closed arm was opened, and the mouse was placed in the starting arm. The previously closed arm was named the « New arm » (N) and the two other arms were named « Others » (O). Parameters - total distance traveled, velocity, the alternation between the arms, entries into the three arms – were measured during 5 min. The short-term spatial memory test was considered successful when the proportion of entries in the new arm was higher than the time spent in the other two arms during the first 2 min of the test.

### Sacrifice and brain tissue preparation

The mice were sacrificed by beheading to prevent any influence of anesthetization (Le Freche et al., 2012). The blood was collected from the neck in heparinized tubes. Brains were removed and one hemisphere was post-fixed for immunohistochemistry in 4% paraformaldehyde fixative in PBS (pH 7.4) for a week at 4°C and transferred to 20% sucrose solution overnight before being frozen. Cortex and hippocampus of the other half of the brain were dissected, disposed in a 1.5 mL isopropylene tube, and then frozen by immersion in an isopropanol solution added with dry ice. Brain tissue was stored at −80°C until used for biochemical analyses. Cortex and hippocampus were added with 200μL of tris-sucrose buffer (TSB) (Tris-HCl 410 mM, pH 7.4 with 10% sucrose) and were sonicated (40 pulses, amplitude 60, 0.5 Hz). A BCA assay was used to determine the protein concentration of each sample.

### Bioavailability assessment

#### Analyte mouse brain extraction

50 mg of the brain were thawed in a safe lock microtube and 500 μL of HCl 1 % and one 5 mm tungsten carbide bead were added. The microtubes were loaded in the TissueLyser II (Qiagen) support plates (24 × 2) at 80°C during 2 × 5 min at 25Hz (between two cycles, 180° plate rotation). The Eppendorf tubes were centrifuged at 12 000 tr/min for 10 min at 4°C. 200 μL of the supernatant were placed on polypropylene tubes and 1800 μL of acetonitrile containing the internal standard (Verapamil 1 nM) at −20°C were added. Each tube was stirred for 30s and placed 1h at −20°C for protein precipitation. The tubes were centrifuged for 10 min at 4°C at 4 000 tr/min.

1800 μL of each tube was removed and transferred in a tube for evaporation in Genevac centrifugal evaporator for 4h30 at 30°C. The residue was dissolved in 200 μL of acetonitrile, vigorously stirred, and evaporated in Genevac centrifugal evaporator for 1h at 30°C. The final residue was dissolved in 90 μL of methanol, vigorously stirred, filtrated, and placed in Matrix tubes for mass spectrometry.

#### Analytical equipment

LC-MS/MS analysis was performed on a UPLC–MS Waters Acquity I-Class coupled to a Xevo TQS Mass Spectrometer (Waters^®^). Instrument control, data acquisition, and processing were made by MassLynx™ software and the reprocessing was carried out using a MassLynx™ sub-software (TargetLynx). The separation was carried out on a Waters^®^ Acquity BEH (C18, 50 × 2.1mm, 1.7μm (40°C)). The injection volume was 1 μL. Elution was performed at a flow rate of 500 μL/min with H2O-ammonium formate 5 mM (pH 3.75) as eluent A and acetonitrile-ammonium formate (5 mM, 5% H2O) as eluent B, employing a 0.1 min plateau with 2% B and a linear gradient from 2% B to 98% B in 1.90 min, followed by a 0.5 min plateau with 98% B. Then, column re-equilibration was performed for 1.5 min. MS analysis was carried out in positive ionization mode using an ion spray voltage of 5000 V. The nebulizer (air) and the curtain (argon) gas flows were set at 0.5 bar. The source temperature and the cone gas flow were set at 150°C and 50 L/h, respectively. The desolvation temperature and desolvation gas flow were set at 600°C and 1200 L/h, respectively. The Multiple Reaction Monitoring (MRM) transitions were monitored with the following values; PEL24-199: 478.40/125.98 and MAGS02-14: 479.40/112.04; Verapamil (Internal Standard): 455.32/165.04. The collision energies were 42 eV (PEL24-199), 46eV (MAGS02-14) and 56eV (Verapamil) for all these transitions.

### Insoluble Tau fraction preparation

70 μL of previously dosed hippocampus samples were added with 130μL of 10% TSB and the samples (crude) were centrifuged at 14000 rpm for 10 min (Centrifuge 5424R, Effendorf). The supernatant (S1) was added with 10% TSB q. s. 600 μL and sonicated before being centrifuged in an ultracentrifuge (Optima TLX rotor TLA-110, Beckman) at 49000 rpm for 1 hour. The supernatant was collected, and the pellet was resuspended with 600 μL of a tris-triton (2%) solution (tris 10 mM pH 7.4, 2% triton) (S2). The samples were sonicated and re-centrifuged at 49000 rpm for 1 hour. The resulting supernatant (S3) was collected, and the pellet (C3) was resuspended in 1 volume of NuPage NuPAGE©LDS 2X sample buffer supplemented with 20% NuPAGE© sample reducing agents (Invitrogen). The regular western-blot protocol was then used and 8 μL of crude, 10 μL of S1, 15 μL of S2 and S3, and 20 μL of C3 were loaded onto the gels. Quantification of protein expression levels was performed using ImageQuantTL Software, and a ratio of insoluble fraction on soluble fraction + insoluble fraction was calculated.

### SDS-PAGE and Western Blot

Hippocampus and cortex samples were prepared in the same concentration in TSB and added 1 volume of NuPAGE LDS 2X sample buffer supplemented with 20% NuPAGE sample reducing agents (Invitrogen) and they were heated 10 min at 70°C. 8μg of proteins per well were loaded onto precast 12% Criterion™ XT Bis-Tris polyacrylamide 26 wells gels (Bio-Rad) and electrophoresis was achieved by applying a tension of 200V during 60 min using a Criterion™ Cell with the NuPAGE MOPS SDS running buffer (1X). Molecular weights calibration was achieved using molecular weight markers (Novex and Magic Marks, Life Technologies). Proteins were transferred to a nitrocellulose membrane of 0.4 μM pore size (G&E Healthcare) using the Criterion blotting system by applying a tension of 100 V for 40 min. Protein transfer and quality were determined by a reversible Ponceau Red coloration (0.2% xylidine Ponceau red and 3% trichloroacetic acid). Membranes were then blocked during 1 hour in 25 mM Tris–HCl pH 8.0, 150 mM NaCl, 0.1% Tween-20 (v/v) (TBS-T) and 5% (w/v) of skimmed milk (TBS-M) or 5% (w/v) of bovine serum albumin (TBS-BSA) depending on the antibody. Membranes were then incubated with primary antibodies overnight at 4°C. Conditions of use of the primary and secondary antibodies are summarized in Table 1. Membranes were rinsed 3 times 10 min with TBS-T and then incubated with secondary antibodies for 45 min at room temperature (RT). The immunoreactive complexes were revealed using a standard ECL detection procedure. Quantifications of protein expression levels were performed with ImageQuantTL Software and the values for the samples were divided by the values of the housekeeping gene (GAPDH). The obtained results for treated samples were divided by the results for control samples to normalize the data.

### Immunohistochemistry and images analysis

Coronal free-floating brain sections of 40 μm were obtained with a cryostat (CM3050 S, Leica). The sections of the hippocampus were selected according to the stereological rules and were stored in PBS (phosphate buffer saline) with 0.2% sodium azide at 4°C. The coronal brain sections were washed with 0.2% Triton X-100 in PBS for permeabilization. Sections were incubated with a 0.3% hydrogen peroxide solution and then blocked with 10% “Mouse on Mouse” Kit serum (ZFO513, Vector Laboratories) for 1 hour before incubation with primary anti-Tau or anti-GFAP/IBA1 antibody overnight at 4°C. Antibodies used in this study are listed in Table 1. After washing in PBS, the sections were incubated with biotinylated anti-mouse or anti-rabbit IgG secondary antibody for 1 hour. Then sections were incubated with the ABC kit (Vector Laboratories) for 2 hours and developed using DAB (Sigma) before being rinsed with physiological serum. Brain sections were mounted on glass slides (Superfrost Plus, ThermoScientific) and dehydrated by sequential baths in 30%, 70%, 95%, and 100% ethanol for 5 min. Then the slides were immersed in toluene for 15 min and fixed with mounting medium (VectaMount Permanent Mounting Medium H-5000, Vector Laboratories) and glass coverslips. Images were acquired using Zeiss Axioscan.Z1 slidescan and quantification of the NFTs-containing neurons was performed by counting the number of events in the CA1 area of the hippocampus, for an average of 3 anteroposterior sections selected according to the Allen mouse brain atlas in 5 mice for each antibody.

### ELISA measurements

The blood samples in heparinized tubes were centrifuged at 10000 rpm for 15 min (Centrifuge 5424R, Eppendorf), and the plasma was collected. Plasma levels of total human Tau protein were measured using ELISA kits (Total Tau ELISA, EuroImmun, EQ6531-9601-L) following the manufacturer’s instructions. Briefly, 100 μL of biotin solution per well were incubated with 25 μL of samples, calibrators, and controls during 3 hours at room temperature. ELISA plate was washed using the washing buffer and 100 μL per well of enzyme conjugate was added for 30 min. The wells were washed again and 100 μL per well of chromogen/substrate were incubated for 30 min covered from light. 100 μL of stop solution per well were added to stop the reaction and absorbance at 450 nm was measured by Multiskan Ascent counter (ThermoLab Systems). The amounts of total Tau were assessed by referring to the standard curve of the manufacturer and expressed as pg/mL.

### Statistics

Results are expressed as means ± SEM. Differences between mean values were determined using the Student’s t-test or a Mann-Whitney U-test using Graphpad Prism Software 8.4.2. p values < 0.05 were considered significant.

## 3 Results

### PEL24-199 treatment restores the short-term memory deficits in a mouse model of Tau pathology

Although NFT are observed in the hippocampus, cognitive impairment appear to be moderate before 6 months in THY-Tau22 mice (Carvalho et al., 2019), and then strengthen at 7 months of age (Sergeant et al., 2019). The associated spatial memory deficits then worsen over time to reach a maximum at 10 months (Schindowski et al., 2006; Van der Jeugd et al., 2013). To investigate whether PEL24-199 or MAGS02-14 compounds, that differ only by one nitrogen atom substitution (Figure 1A), had or not a similar *in vivo* activity, global behavioral and short-term spatial memory were assessed at 7 months of age, a stage at which THY-Tau22 mice already exhibit spatial memory impairments (Sergeant et al., 2019) and ongoing Tau pathology development, following 6 weeks, i.e. in a curative paradigm (Figure 1B). The anxiety measured using the Elevated Plus Maze test showed no significant impact of PEL24-199 and MAGS02-14 treatments on velocity, average distance moved, or percentage of time spent in the closed or the open arms for either WT or THY-Tau22 mice (*p = 0.53*, Supplementary figure 1). Thus, the treatments did not significantly affect the locomotion and basal anxiety behavior of both WT and THY-Tau22 mice, which overall means that the short-term spatial memory effect of our molecules is not influenced by locomotion or anxiety changes. In the Y-maze task, which evaluate short-term spatial memory, 7 months WT mice treated at 1 mg/kg with MAGS02-14 spent less time in the new arm compared to the untreated WT mice (Figure 1C). At the same dose, PEL24-199 did not alter the performance of the WT mice. As expected, at 7 months of age, THY-Tau22 mice exhibited short-term spatial memory impairment with an absence of preference for the new arm. MAGS02-14 treatment had no significant effect on the spatial memory of THY-Tau22 mice. In contrast, PEL24-199 mitigated memory impairments of THY-Tau22 mice.

**Figure 1:**
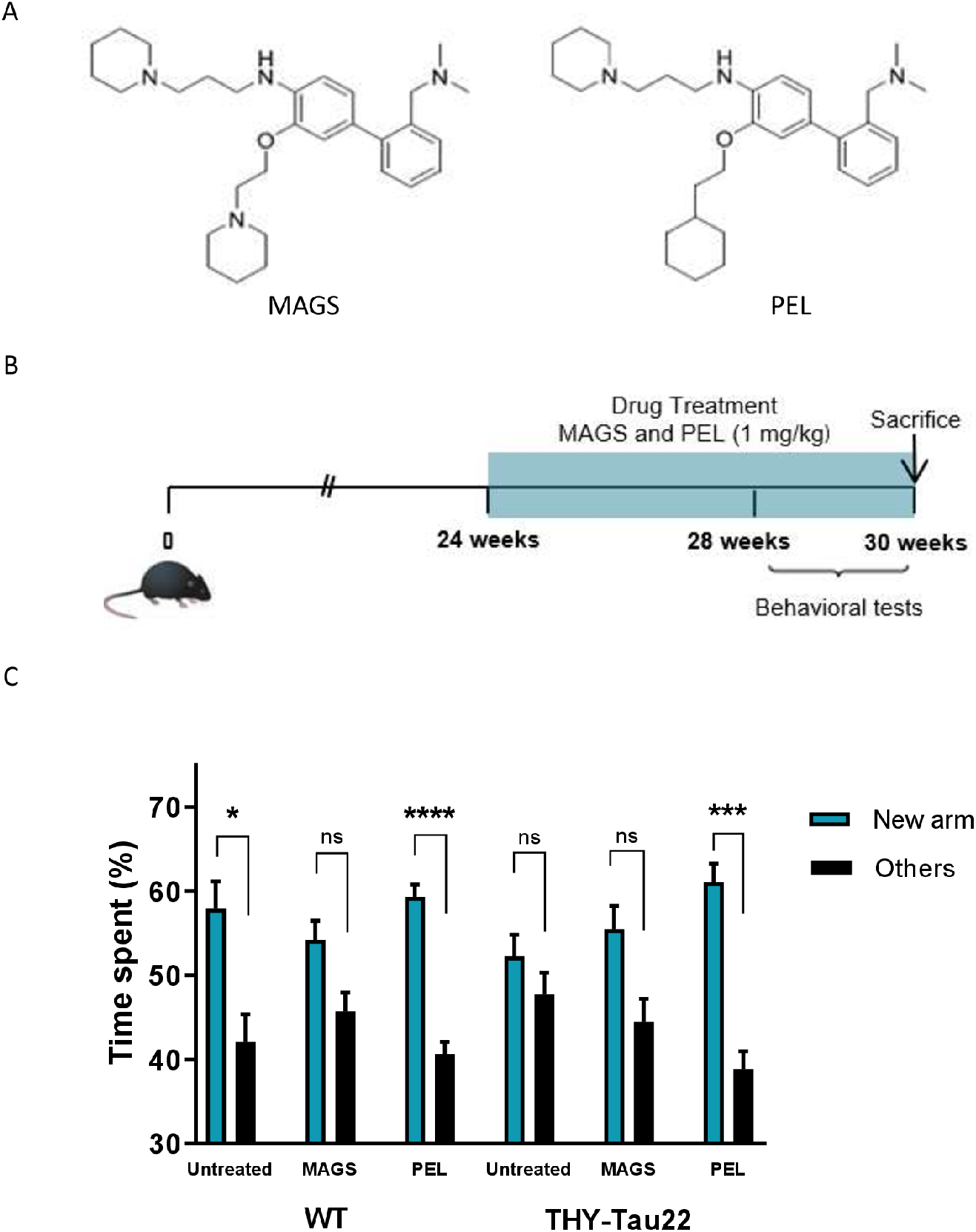
PEL24-199 restores short-term memory in a mouse model of Tau pathology. **A.** Chemical structure of MAGS02-14 (referred to as MAG-S) in and PEL24-199 (referred to as PEL) compounds, differing by one nitrogen atom. **B.** Temporal representation of THY-Tau22 mice treatment. The mice were treated from 24 weeks to 30 weeks with 1mg/kg of PEL24-199 or MAGS02-14. **C.** PEL24-199 treatment of THY-Tau22 with 1mg/kg improved short-term memory in the Y-maze task compared to untreated animals. Animals were placed in a three-arm maze surrounded by spatial clues and one arm was closed during the learning phase of 5 min. The animals were returned to the home cage for the retention phase of 2 min. Then the third arm was opened, and the time spent in each arm by the animals was recorded for 5 min. Histograms represent the means±SEM (n=12 animals per condition), *: p <0.05, **: p <0.01, ***: p <0.001 ****: p <0.0001 using Mann-Whitney statistical test. Abbreviations: SEM, standard error of the mean.

### PEL24-199 decreases hyperphosphorylated Tau in mice brain extracts

Cognitive impairment is associated with the progression of Tau pathology in the hippocampus and the cortex of THY-Tau22 mice (Van der Jeugd et al., 2013). Any memory impairment could therefore relate to a modification of Tau pathology and Tau phosphorylation status. We therefore assessed hippocampal and cortical Tau expression and phosphorylation with antibodies raised against N- and C-terminus of Tau proteins, phospho-sites that are known to be hyperphosphorylated in AD (_S_396 and _S_262, Augustinack et al., 2002) as well as pathological epitopes which are only detected when neurofibrillary processes are present (_T_212/_S_214 and _S_422) (Figure 2A). Treatment with PEL24-199 or MAGS02-14 did not affect the expression of total Tau protein either in the hippocampus or in the cortex of THY-Tau22 mice (Figure 2B). MAGS02-14 treatment showed no significant effect on Tau phosphorylation at this concentration, either on physiologic or pathological epitopes (Figure 2C). Interestingly, PEL24-199 decreases by half the _S_396 and _S_262 epitopes in the cortex, and decreases significantly pathological epitopes _T_212/_S_214 and _S_422 in this structure. A decrease in the staining of those antibodies is indicative of a reduction of the neurofibrillary degenerating process. PEL24-199 also reduced the phosphorylation of the _S_262 and _S_396 (*p = 0.0625*) epitopes and strongly decreases the phosphorylation on the _T_212/_S_214 (*p = 0.0625*) and _S_422 sites in the hippocampus of THY-Tau 22 treated mice (Figure 2C). Levels of unphosphorylated Tau at 198-204 amino acid sequence (numbering according to the longest 441 human brain Tau isoform) did not change under PEL24-199 treatment. Tau phosphorylation is under the control of phosphatase and kinases, the level of expression of the principal Tau serine/threonine phosphatase PP2A (Liu et al., 2005) was then investigated. The results show that MAGS02-14 had no impact on PP2A levels neither in the cortex nor in the hippocampus (Figure 2 D, E). Interestingly, the levels of PP2A in the cortex were not affected by treatment with PEL24-199 (Figure 2D); however, there was a sharp increase of PP2A in the hippocampus of THY-Tau22 mice treated with PEL24-199 (Figure 2E).

**Figure 2:**
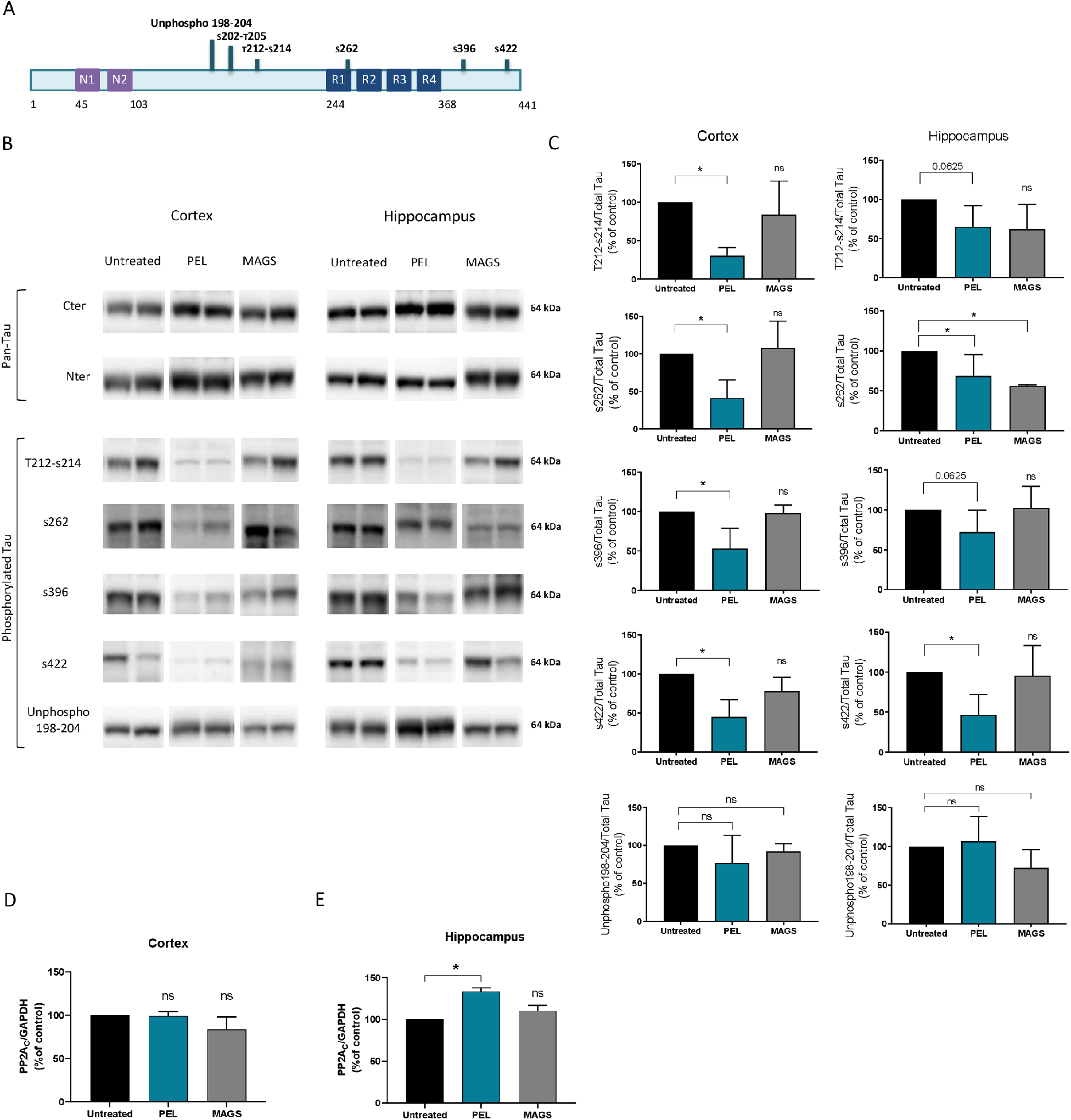
PEL24-199 decreases Tau hyperphosphorylation and increases PP2A expression in the brain of THY-Tau22 mice. **A**. Representation of the phosphorylation sites studied within the Tau protein. **B**. Western blotting of brain lysates from Thy-Tau22 mice (n=6) or Thy-Tau22 mice treated with 1 mg/kg of PEL24-199 (n=6) was used to assess the phosphorylation of tau at serine 212-214 (pSer 212-214), serine 262 (pSer262), serine 396 (pSer396), serine 422 (pSer422), unphosphorylated Tau consisting of residues 198–205 (Tau-1) and overall Tau expression (pan-Tau antibodies against the N-terminus (Nter) and C-terminus (Cter)). **C.** Western-blot quantification of Tau phosphorylation sites in the hippocampus and the cortex, expressed as a percentage of the control condition. **D, E.** Western-blot quantification of PP2A phosphatase expression in cortex and hippocampus. GAPDH staining was used as a marker and loading control. Histograms represent the means±SEM. The mean difference was statistically analyzed using a Mann-Whitney test: *: p <0.05.

### PEL24-199 decreases detergent-resistant phospho-Tau in mice hippocampus

In THY-Tau22 mice, Tau aggregation is associated with increased insolubility of Tau (Schindowski et al., 2006). We then assessed the soluble and insoluble Tau fractions from THY-Tau22 mice treated with either MAGS02-14 or PEL24-199. While MAGS02-14 treatment exhibits no significant decrease of the insoluble Tau fraction (Supplementary figure 2) when compared to THY-Tau22 untreated animals, insoluble Tau fraction (C3) was significantly decreased in mice treated with PEL24-199 (Figure 3A), as visualized with the Tau-Nter as well as with the Tau _S_396 antibodies (Figure 3C, D). To a lesser extent, PEL24-199 treatment non-significantly increased the unphosphorylated 198-204 antibody in Tau insoluble fraction (Figure 3E). While MAGS02-14 had no effect on the cognitive function nor on Tau phosphorylation and Tau insolubility, PEL24-199 associates a restored short-term memory, a reduced Tau phosphorylation with a decreased Tau and phospho-Tau insolubility in THY-Tau22 treated animals.

**Figure 3:**
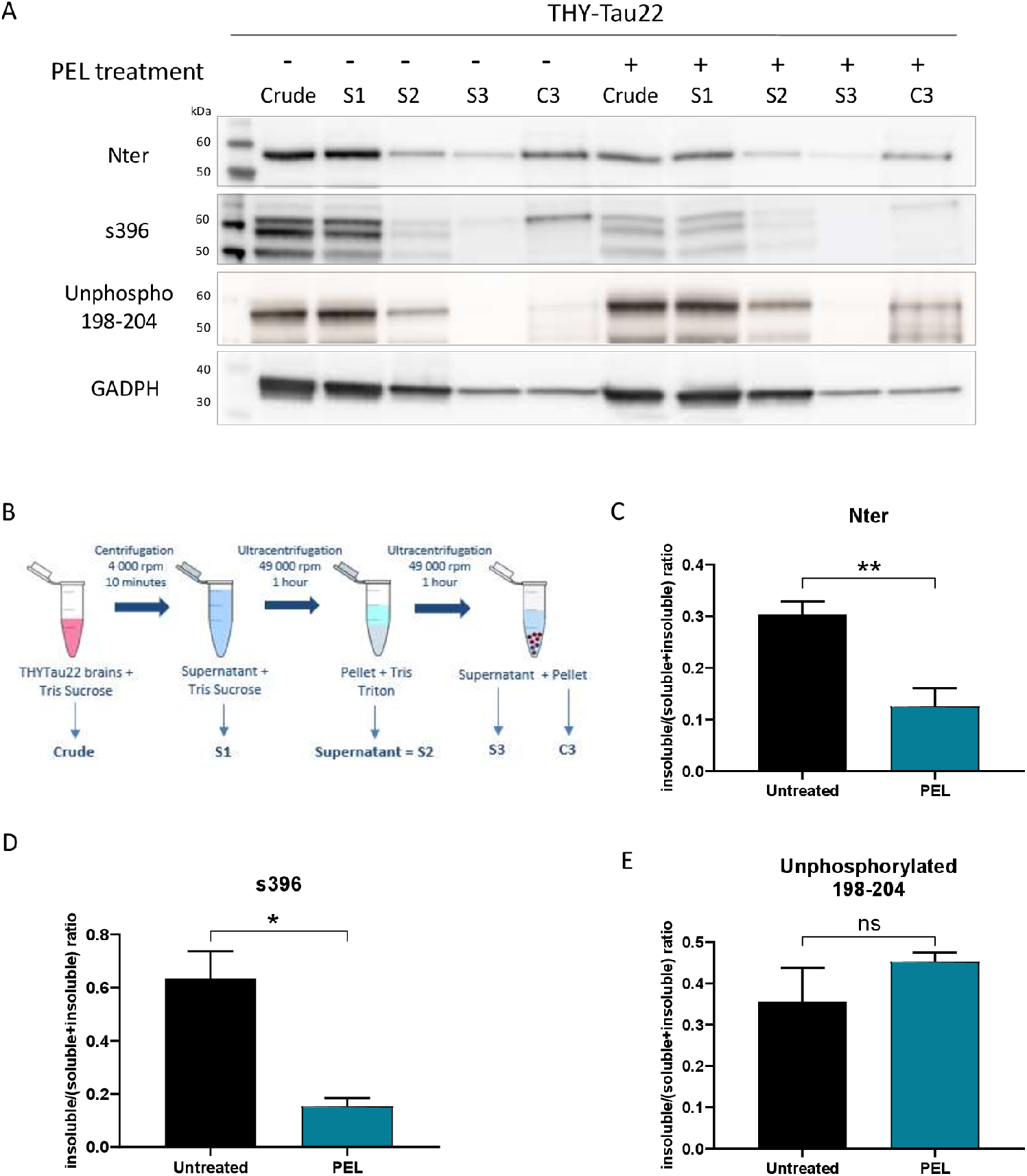
PEL24-199 decreases detergent-resistant phosphorylated Tau in THY-Tau22 hippocampus. **A.** Western-blot analysis of hippocampus lysates of THY-Tau22 (n= 6) and THY-Tau22 treated with PEL24-199 (n=6) with antibodies Nter, pSer396, and Unphosphorylated tau 198-204. **B.** Schematic representation of fractionation steps. The samples in TSB (crude) are centrifuged and the supernatant is added with 10% TSB (S1) and sonicated before being centrifuged. The supernatant is collected (S2) and the pellet is resuspended in a tris-triton solution. The samples are sonicated and re-centrifuged. The obtained supernatant (S3) is collected and the pellet is resuspended in LDS buffer (C3). **C, D, E.** Western-blot quantification of Nter, pSer396 and Unphosphorylated tau 198-204 antibodies in the hippocampus lysates of THY-Tau22 and THY-Tau22 treated with PEL24-199 (n=6). Results are represented as histograms of the mean ± SEM, expressed as a percentage of the total fraction. The mean difference was statistically analyzed using a Mann-Whitney test: *: p <0.05.

### PEL24-199 reduces NFTs and astrogliosis in the hippocampus of THY-Tau22 treated mice

THY-Tau22 mice exhibit neurofibrillary tangle-like inclusions with rare ghost tangles and PHF-like filaments, as well as mild astrogliosis (Schindowski et al., 2006). To further assess the modulatory effect of our compounds, the burden of NFT in the hippocampal CA1 was investigated by immunohistochemistry using antibodies against hyperphosphorylated Tau epitopes _S_202/_T_205 and _S_396/404 and pathological Tau phospho-sites _T_212/_S_214 (Figures 4A). MAGS02-14 did not show any reduction of neurofibrillary degeneration bundle (Supplementary figure 3). In THY-Tau22 animals treated with PEL24-199, _S_202/_T_205 and _S_396/_S_404 epitopes were significantly reduced, underlying a decrease of Tau hyperphosphorylation in neurons of the hippocampus (Figure 4B). The pathological _T_212/_S_214 epitope was reduced although not significantly (*p = 0.09*), showing a tendency for PEL24-199 to also reduce Tau pathology. Overall, these results show a reduction of the number of NFT containing neurons in THY-Tau22 mice treated with PEL24-199. Astroglial activation has been shown to be activated with the development of Tau pathology (Laurent et al., 2017; Laurent et al., 2018) and presumably favors the development of Tau pathology in a detrimental circle (Laurent et al., 2018; Ising et al., 2019). We therefore further investigated the impact of PEL24-199 treatment on astrocytes activation. Our data showed that this molecule lowered GFAP staining using immunohistochemistry (Figure 4 C, D). PEL24-199 is therefore able to decrease astroglial activation in the hippocampus of THY-Tau22 mice.

**Figure 4:**
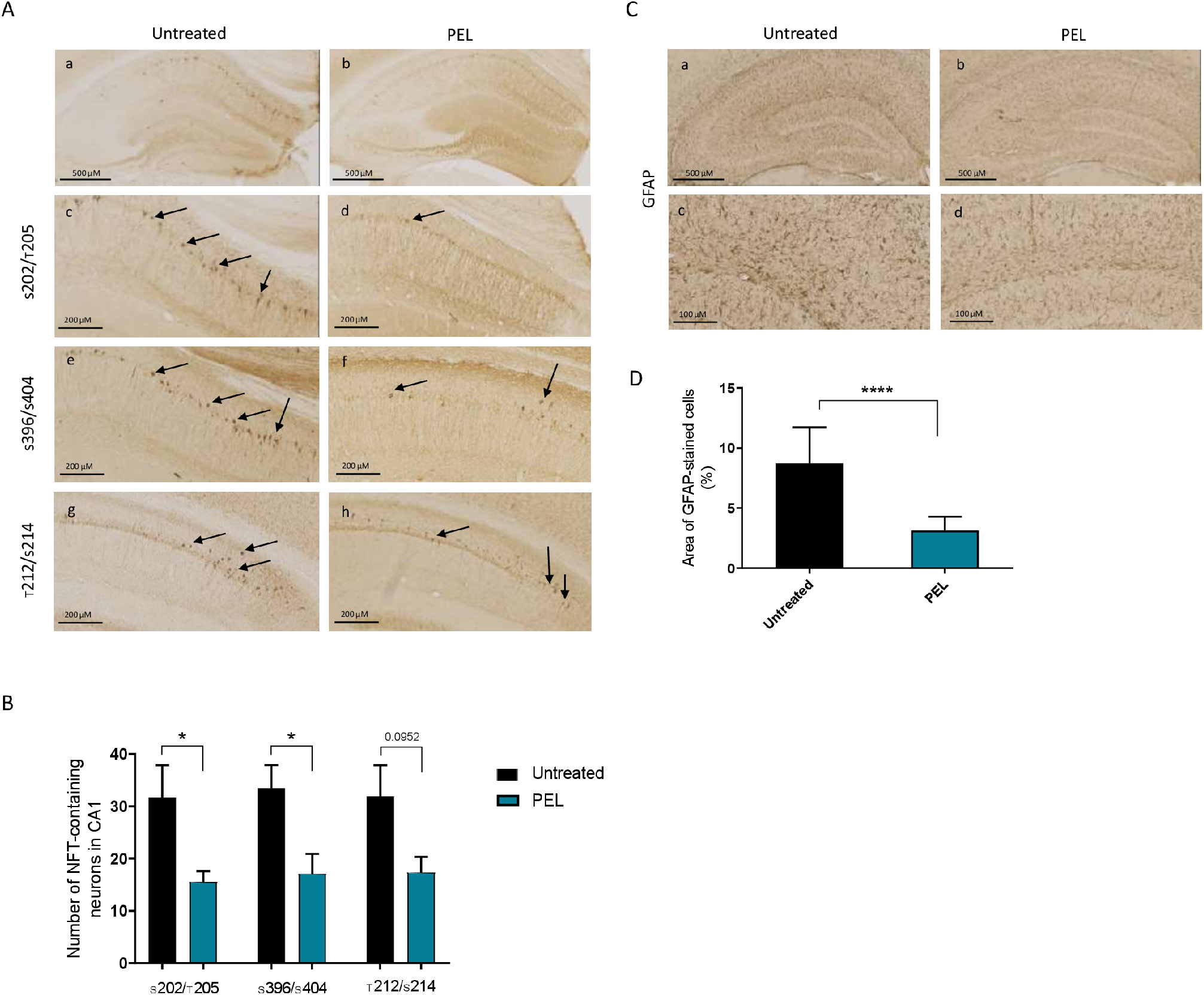
PEL24-199 reduces Tau pathology and astrogliosis in the hippocampus of Thy-Tau22 treated mice. **A.** Histological analysis of neurofibrillary degeneration using phospho-Tau antibodies against phospho-epitopes (_S_202/_T_205 and _S_396/404) or pathological epitopes (_T_212/_S_214) in THY-Tau22 mice and THY-Tau22 treated with PEL24-199. **B**. The number of NFT-labeled neurons calculated from 3 brain slices in each animal (n=5) showed significantly reduced _S_202/_T_205 and _S_396/404 labeling with PEL24-199 treatment and a reduction for _T_212/_S_214 labeling although not significant (Mean± SEM, *: p <0.05, **: p < 0.01, Mann-Whitney t-test). **C.** Histological analysis of astrocytic activation performed on hippocampus sections of THY-Tau22 mice and THY-Tau22 mice treated with PEL24-199. Astrocytic activation is marked thanks to the anti-GFAP antibody. **D.** Quantification of the anti-GFAP antibody. The total volume of labeling (calculated in percentage of the total area of the hippocampus) calculated from three brain slices in n=5 animals showed that PEL24-199 treatment decreased the number of reactive astrocytes (****: p <0.0001, Student’s t-test. 3 brain slices in each animal (n=5)).

## 4 Discussion

In the present study, we show that the β-secretase non-competitive inhibitor compound PEL24-199 represses the Tau pathology, reduces the astrogliosis, and improves short-term spatial memory in the THY-Tau22 transgenic model of hippocampal neurodegeneration. These results, therefore suggest that this APP metabolism regulatory PEL24-199 compound mitigates the Tau pathology *in vivo*. This effect is observed in a curative paradigm and results are in line with previous studies where molecules derived from the same pharmacophore were effective in a preventive paradigm against both amyloid and Tau pathologies (Sergeant et al., 2019).

A compound containing the same pharmacophore but fused with a tacrine moiety, RPEL, was shown to reduce both the amyloid pathology in the APPxPS1 transgenic animals and tau pathology in the THY-Tau22 hippocampal neurofibrillary degeneration model (Sergeant et al., 2019). These effects were also associated with a cognitive improvement although in a preventive paradigm since animals were treated starting from the age of 3 months and before the appearance of lesions in both transgenic models. Taking herein advantage of the structure-activity relationship, two compounds differing by a single nitrogen atom and sharing a β-secretase non-competitive inhibitory effect but having a lysosomotropic activity for MAGS02-14 only were compared (Gay et al., 2018). This lysosomotropic activity is common to several compounds that were originally derived from chloroquine (Melnyk et al., 2015). Through the alkalization of intravesicular pH, the lysosomotropic activity of compounds inhibits the β-secretase pH-dependent activity and represses the autophagic flux (Schrader-Fischer and Paganetti s. d.; Tam et al., 2014). Dosage of MAGS02-14 in the brain tissue showed an accumulation when compared to PEL24-199 (Supplementary figure 4). This accumulation could contribute to the inefficacy of MAGS02-14 which accumulation in brain tissue could be deleterious. Modulation of the g-secretase which is also routed to the early endosome together with the β-secretase is likely not contributing to the observed effect of our compounds since Notch1 g-secretase processing is not modified by RPEL, MAGS02-14, or PEL24-199 (Gay et al., 2018; Sergeant et al., 2019). Together, our results suggest that the lysosomotropic activity is dispensable for the in *vivo* activity of our compounds whereas the β-secretase non-competitive inhibitory activity is more likely essential.

Although a direct relationship between β-secretase aspartyl proteases BACE1 or BACE2 and Tau protein expression has not yet been described, a growing body of evidence suggests an interplay between Tau protein and the β-secretase processing of APP. β-secretase inhibitors or g-secretase modulators were shown to reduce Tau protein expression in control neurons derived from human stemcell-derived excitatory cortical neurons (Moore et al., 2015). Following PEL24-199 treatment of THY-Tau22 mice, Tau phosphorylation was reduced at hyperphosphorylated sites and pathological phosphosites. Moreover, the insoluble fraction of Tau was reduced as well as the number of neurofibrillary tangles. Notably, the decrease of Tau phosphorylation was not followed by an increase in Tau plasmatic clearance (Supplementary figure 5), suggesting that the positive effects observed with PEL treatment on the decrease of Tau pathology are to a change with the plasma clearance of Tau. Change of Tau phosphorylation could be due to the modification of PP2A expression like hyperphosphorylation of Tau Ser202/Thr205 is negatively regulated by PP2A activity (Kins et al., 2003) or increased activation of PP2A contributes to the restoration of cognitive functions in THY-Tau22 mice, also in a curative paradigm (Ahmed et al., 2020). PP2A is inhibited in AD and suggested to contribute to the hyperphosphorylation of Tau and the regulation of APP metabolism (Taleski et al., 2021). PP2A expression is increased in THY-Tau22 mice treated with PEL24-199 but not those treated with MAGS02-14, first showing the specific effect of PEL24-199, and secondly, we can assume a relationship between the reduction of Tau phosphorylation and increase expression of PP2A. In PEL24-199-treated mice insoluble Tau fraction also decreased indicating that the proportion of aggregated Tau is diminished resulting either from the lowering of existing neurofibrillary degenerating processes or the inhibition of this process or both. These results are strengthened by the significant lowering of the number of neurofibrillary degenerating neurons in the brain of PEL24-199-treated animals. We therefore demonstrated that PEL24-199, like RPEL (Sergeant et al., 2019), can decrease Tau pathology *in vivo* by reducing the number of neurofibrillary tangles present in the hippocampus. Together, these results demonstrate a reduction of the neurofibrillary degenerating process in THY-Tau22 treated mice when compared to untreated animals and therefore PEL24-199 compound reduces Tau pathology in a curative paradigm together with the recovery of the short-term spatial memory. Our results are in line with the paper of Moore et al (Moore et al., 2015), in which they have shown that manipulating APP metabolism by a β-secretase inhibition results in a specific decrease in Tau protein levels, demonstrating that APP metabolism regulates Tau proteostasis. Our data also show that modulating the metabolism of APP with small molecules can affect not only Tau protein levels but also the neurofibrillary degenerating process and *in fine* improve cognitive functions.

Few studies involving β-secretase inhibitors have shown to reverse or attenuate behavioral and memory deficits in transgenic mouse models of AD (Imbimbo and Watling, 2019). Research of therapeutics for neurodegenerative diseases has put forward several small molecules as candidates either targeting Aβ or Tau lesions (Morimoto et al., 2013; Lecoutey et al., 2014; Yahiaoui et al., 2016), including autophagy modulators (Silva et al., 2020), but to our knowledge, none of them are acting on both amyloid and Tau pathological processes. BACE1 and BACE2 were shown to degrade Aβ peptides besides being the protease necessary to produce Aβ peptides (Abdul-Hay et al., 2012). Thus, current inhibitors may also affect Aβ degradation through uncomplete repression of the aspartyl protease activity of BACE proteases (Liebsch et al., 2019). As PEL24-199 is not a direct inhibitor of BACE1, this compound may modulate differently the APP metabolism and therefore potentially preclude the detrimental effect of pure β-secretase inhibitors.

Astrogliosis is an inflammatory response that potentiates the progression of neurodegenerative diseases and is considered a potential therapeutic target (Phillips et al., 2014; Chung et al., 2015). Astrocytes have a discrete regulatory function of synapses and neuronal plasticity and for instance, specific reduction of Connexin 43 in astrocytes reduces the memory impairment in APPxPS1 mice (Ren et al., 2018). Levels of GFAP-reactive astrocytes are closely associated with dementia in AD (Perez-Nievas et al., 2013). More recently, senescent astrocytes accumulation was shown to promote the formation of hyperphosphorylated tau aggregates while reducing those senescent astrocytes prevents PS19 Tau transgenic mice from cognitive decline and reduced Tau pathology (Bussian et al., 2018), showing a close interplay between the Tau pathology and reactive astrogliosis. Herein, we showed that GFAP-positive reactive astrocytes were reduced in THY-Tau22 mice treated with PEL24-199 when compared to untreated mice. This GFAP-reactive astrocyte reduction could either result from PEL24-199 direct effect on astrocyte or indirectly related to the reduction of the Tau pathology. Suppression of Tau expression in the double APP/PS1 × rTg4510 transgenic as well as in rTg4510 transgenic model of Tau pathology reduced the burden of NFT, the astrogliosis even in greater proportion in the single rTg4510 (DeVos et al., 2018). These results taken together could suggest that the reduced astrogliosis in the THY-Tau22 treated with PEL24-199 could be attributable in part to a direct effect of PEL24-199. This reduced astrogliosis may also contribute to the cognitive improvement observed in PEL24-199 treated animals.

In the present study, we showed that PEL24-199, but not MAGS02-14, leads to a restoration of cognitive functions but also a reduction of the Tau pathology and associated astrogliosis in the Tau pathology transgenic model THY-Tau22. The effect of our molecule relies on a modification of APP processing through a non-competitive β-secretase modulation effect and where the lysosomotropic activity is dispensable. Thus, PEL24-199 treatment in curative paradigm reduce the Tau pathology, the astrogliosis and restore the short-term memory. Together, these results indicate that we have a molecule functional on APP metabolism (Gay et al., 2018, Tautou et al., unpublished data) but also on Tau pathology. Further investigations will be necessary to elucidate the precise molecular mechanism of action of these molecules that are effective against both amyloid and Tau pathology.

## Supporting information

Tautou et al supplementary material and results

## 6 Author Contributions

MT contributed to the data acquisition, analysis, interpretation, and wrote the manuscript. SE contributed to the data acquisition and analysis. FD and P-EL provided study materials. CL contributed to the analysis. DB and LB contributed to the design of the study. PM and NS contributed to the design of the study, interpretation, and writing of the manuscript. All authors read and approved the submitted manuscript version.

## 8 Acknowledgments

We thank the BICeL (Imagerie Cellulaire et Tissulaire) platform, the IVF (Imagerie du Vivant et Fonctions - Plateau d’explorations fonctionnelles) platform, Genotyping platform and Lille ADME platform (U1177), particularly Sébastien Carrier and Catherine Piveteau. The 300 MHz NMR facilities were funded by the Région Nord Pas-de-Calais (France), the Ministère de la Jeunesse, de l’Education Nationale et de la Recherche (MJENR) and the Fonds Européens de Développement Régional (FEDER). We would like to thank Dihia Amara and Katia Arab for their technical assistance and the SPQI Company for providing phospho-Tau antibodies.

