## Supplementary material for "A β-secretase modulator decreases Tau pathology and preserves short-term memory in a mouse model of neurofibrillary degeneration": Tautou et al supplementary material and results

### 1 Supplementary Figures

| Name | Species | Dilution | Saturation | Provider & Reference |
|---|---|---|---|---|
| **Western Blot – Primary antibodies** | | | | |
| **Tau Nter** | Rabbit | 1 : 5000 | TNT/5% milk | Homemade 12-41 Rabbit 1242075 |
| **Tau Cter** | Rabbit | 1 : 10 000 | TNT/5% milk | Homemade 993-S2 Y16L-2P [427-441] |
| **S262** | Rabbit | 1 : 5000 | TNT/5% BSA | Invitrogen 44750G |
| **S396** | Rabbit | 1 : 10 000 | TNT/5% BSA | Invitrogen 44752G |
| **S422** | Mouse | 1 : 1000 | TNT/5% BSA | 4BDX 1501 |
| **T212-S214 (AT100)** | Mouse | 1 : 1000 | TNT, no saturation | Invitrogen MN1060 |
| **Unphospho 198-204** | Mouse | 1 : 2000 | TNT/5% milk | Millipore MAB3420 |
| **GAPDH** | Rabbit | 1 : 50 000 | TNT/5% milk | Sigma G9545 |
| **PP2A C subunit** | Rabbit | 1 : 1000 | TNT/5% milk | Sigma 07-324 |
| **Western Blot – Secondary antibodies** | | | | |
| **Goat Anti-Rabbit** | Goat | 1 : 5000 | | Vector PI1000 |
| **Goat anti-Mouse** | Goat | 1 : 50 000 | | Merck Millipore AP200P |
| **Immunohistochemistry – Primary antibodies** | | | | |
| **S202-T205 (AT8)** | Mouse | 1 : 500 | | Thermo scientific MN1020 |
| **T212-S214 (AT100)** | Mouse | 1 : 500 | | Invitrogen MN1060 |
| **S396-S404** | Mouse | 1 : 4 | | Made in the lab |
| **GFAP** | Rabbit | 1 : 1000 | | Dako, Z0334 |
| **Immunohistochemistry – Secondary antibodies** | | | | |
| **Goat anti-Mouse IgG biotinylated** | Goat | 1 : 400 | | Vector ZF0805 |
| **Goat anti-Rabbit IgG biotinylated** | Goat | 1 : 400 | | Vector BA-1000 |

**Supplementary Table 1. List of antibodies used for immunohistochemistry and western-blot analyses.**

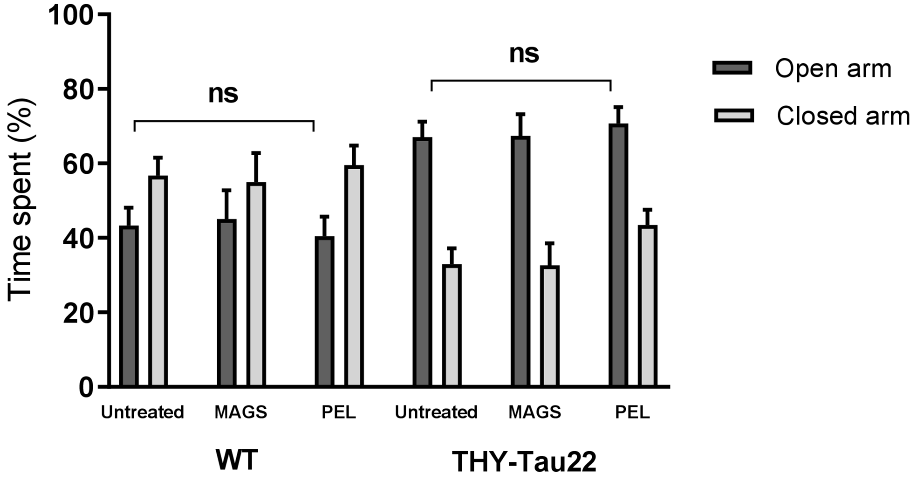


**Supplementary Figure 2. PEL24-199 and MAGS02-14 had no influence on THY-Tau22 mice anxiety.** PEL24-199 or MAGS02-14 treatment of THY-Tau22 with 1mg/kg did not influence anxiety in the Elevated-Plus-Maze task compared to untreated animals. Mice movements were recorded for 5 minutes in a plus-shaped maze consisting of two 10-cm-wide open arms and two 10-cm-wide enclosed arms elevated at 50 cm above the floor. Histograms represent the means±SEM (n=12 animals per condition). The mean difference was statistically analyzed using a Mann-Whitney test.

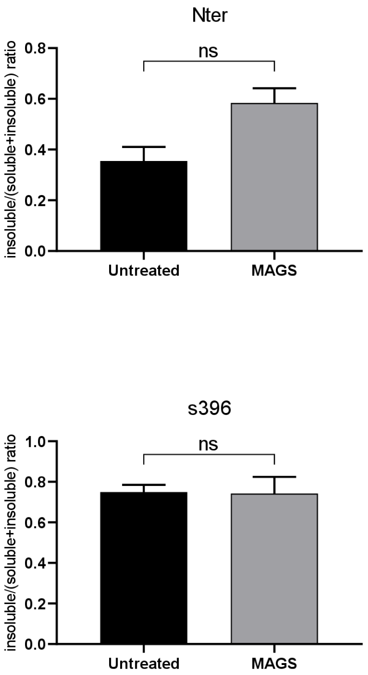


**Supplementary Figure 2. 1 mg/kg of MAGS02-14 did not affect detergent-resistant phosphorylated tau in THY-Tau22 hippocampus**. Western-blot quantification of Nter, pSer396 and Unphosphorylated tau 198-204 antibodies in the hippocampus lysates of THY-Tau22 and THY-Tau22 treated with MAGS02-14. Results are represented as histograms of the mean ± SEM, expressed as a ratio of insoluble fraction on soluble fraction + insoluble fraction. The mean difference was statistically analyzed using a Mann-Whitney test (n = 6 animals for each condition).

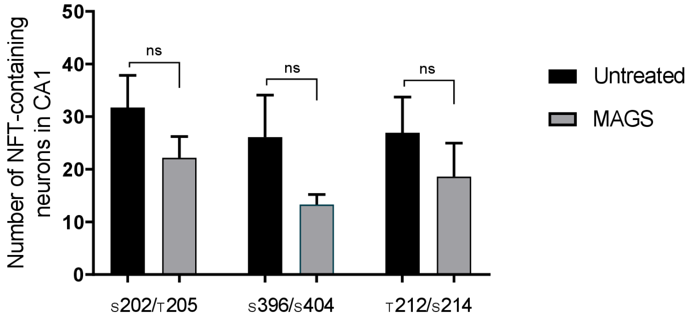


**Supplementary Figure 3. 1 mg/kg of MAGS02-14 had no effect on tau pathology in the hippocampus of Thy-Tau22 mice.** The number of NFT-labeled neurons calculated from 3 brain slices in each animal (n=5) showed no significant modification of p202/205, p396/404 or p212/214 labeling with MAGS02-14 treatment (Mean± SEM, Mann-Whitney t-test).

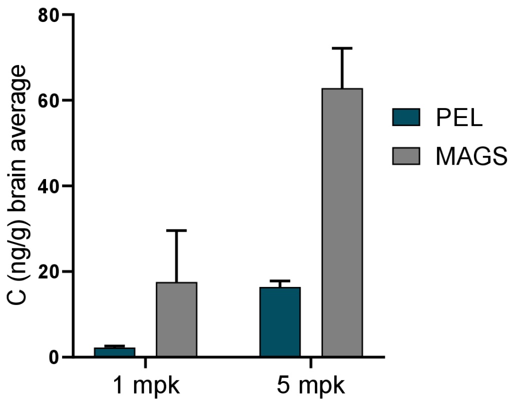


**Supplementary Figure 4. Mass spectrometry quantification of PEL24-199 and MAGS02-14 in WT mice brain**. PEL24-199 and MAGS02-14 were extracted from brains of WT mice treated with 1 mg/kg (mpk) or 5mg/kg and quantified thanks to UPLC- MS/MS TQS procedure, showing the presence and the accumulation of the both compounds in mice brain. The analytical method was validated with specificity, linearity, fidelity and accuracy criteria. Histograms represent the means±SEM (n = 4 brains for each condition).

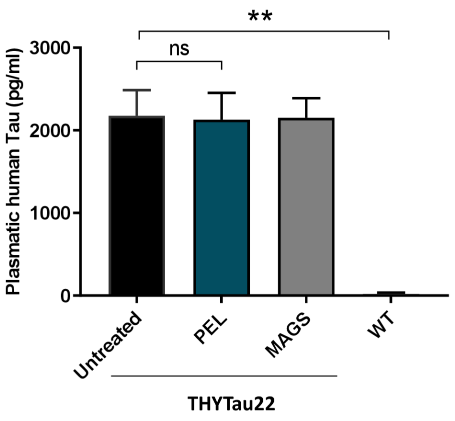


**Supplementary Figure 5. Quantitative ELISA measurements of plasmatic Tau levels of WT mice and untreated, MAGS02-14-treated or PEL24-199-treated THY-Tau22 mice.** Histogram represent the means±SEM (n=7 animals per condition). The mean difference was statistically analyzed using a Mann-Whitney test: ****: $p < 0.0001$.
